# Reduced PTPRD expression differentially alters brain phosphotyrosine phosphoproteomic profiles of 2 and 12 month-old mice

**DOI:** 10.1101/2021.03.02.433536

**Authors:** George R Uhl, Ian M Henderson, Maria Martinez, Matthew P Stokes

## Abstract

The receptor type protein tyrosine phosphatase PTPRD is implicated in maturation of synapses of expressing neurons, vulnerability to addictions, reward from addictive substances, vulnerability to restless leg syndrome and densities of neurofibrillary pathology in Alzheimer’s disease brains by a variety of evidence. However, PTPRD’s physiological substrates and adaptations to differences in levels of PTPRD expression in brains of young and aging animals have not been explored in depth. We report phosphoproteomic studies of brains of young and aged mice with different levels of PTPRD expression, gene ontology studies of genes identified in this way and validation of several candidate PTPRD substrates with *in vitro* assays using recombinant PTPRD phosphatase. PTPRD is well positioned to modulate the extent of phosphorylation of phosphotyrosine phosphoprotein substrates, including those involved in synaptic maturation and adaptation.

## Introduction

The receptor type protein tyrosine phosphatase PTPRD is one of the most highly-expressed neuronal protein tyrosine phosphatases (1, 2). PTPRD binds to extracellular ligands including interleukin 1 receptor like and accessory (IL1RL1 and IL1RAPL1), Slittrk, SALM and/or NGL proteins in ways that (likely) alter activities of its intracellular protein tyrosine phosphatase and thus contribute to the maturation of synapses of PTPRD-expressing neurons (1, 3).

Common variation in the human PTPRD gene intron 9 - 11 region exerts sizable “oligogenic” influences on densities of neurofibrillary pathology in Alzheimer’s disease brains, vulnerability to restless leg syndrome (RLS) and levels of PTPRD mRNA in human postmortem cerebral cortex (4–6). There are more modest, polygenic influences of common variation in this gene on vulnerability to development of a substance use disorder, abilities to quit smoking, ability to stop use of opiates and reward from psychostimulants (1, 7, 8). A rare human PTPRD knockout provides substantial intellectual disability (9).

Mice with constitutively-reduced PTPRD expression display reduced reward from stimulants, supporting polygenic associations with addiction phenotypes (5, 10). They display disrupted sleep onset, supporting associations with RLS (5, 10). Homozygous knockouts, but not heterozygotes, display substantial mnemonic deficits (5, 11) in ways that fit with observations in humans. Our initial observations support increased activating tyrosine phosphorylation of the tau kinases GSK3β and GSK3α and greater age-related accumulation of pS202/pT205/pS208 phosphotau immunoreactivity in brains of mice with reduced PTPRD expression (12). Attention to roles of altered PTPRD in aging could thus add insight into potential biochemical bases for reported oligogenic associations between PTPRD variants and the age-related neurofibrillary pathology found in Alzheimer’s disease brains (4). Phosphotyrosine phosphoproteomic studies can identify greater or less abundance of phosphotyrosine phosphoproteins in brains of mice with reduced PTPRD expression. Simplistically, phosphotyrosine phosphoproteins that display increased expression in knockouts and co-expression with PTPRD are candidate PTPRD substrates. Proteins displaying decreased tyrosine phosphorylation in PTPRD knockouts or those that are display increased phosphorylation but are not co-expressed are candidates to provide adaptations to reduced PTPRD expression. Each of these categories is of interest for interpreting the combined human and mouse data relevant to addiction and RLS noted above. When we compare these patterns in young *vs* aged mice with constitutively-altered PTPRD expression, we can assess age-related changes in both candidate PTPRD substrates and in adaptations to altered PTPRD expression. These aging findings could contribute to understanding PTPRD’s associations with Alzheimer’s disease neurofibrillary pathology.

We now report assessment of the abundance of tyrosine phosphorylated brain proteins in brains of 2 and 12 month old wildtype mice and littermates with reduced PTPRD expression. We compare assessments from a complete dataset to those from prior and concurrent work with relevant *c. elegans* and mouse phosphotyrosine phosphoproteomics (13, 14) and with data from a second independent partial dataset from our laboratory. We confirm that several of the candidate PTPRD phosphotyrosine substrates identified by this work are avidly dephosphorylated by recombinant PTPRD phosphatase *in vivo.* We discuss the implications and limitations of these data for understanding PTPRD’s physiological activities and adaptations to its modulation.

## Materials and Methods

### Mice and brains

Mice with reduced PTPRD expression and wildtype littermates were bred from heterozygote x heterozygote crosses as described from mice initially generously supplied by Uetani and colleagues (5, 10, 11). Mice were housed in the NMVAHCS animal facility, fed moistened food on cage floors until weaning and genotyped as described (10). One mouse required sacrifice due to dental issues at about 6 mos of age. Mice were sacrificed by rapid cervical dislocation at about 2 or 12 months of age, brains rapidly removed and split by a midsagittal razor blade cut. Our main dataset comes from analyses of proteins from half brains (n = 4/genotype/age group except n = 3 for 12-month old WT) that were rapidly frozen on aluminum foil placed on dry ice blocks and then rapidly covered by dry ice powder and maintained at −80°C until analyses. A second “partial” dataset was obtained from brains placed into polypropylene tubes that were then floated onto liquid nitrogen in ways that provided differences in time to freezing. All procedures were approved by the NMVAHCS Animal care and use committee.

### Phosphotyrosine phosphoproteomic analyses

We used mass spectrographic analyses and quantitation of immunoprecipitated tryptic phosphotyrosine phosphopeptides extracted from frozen mouse half brains as described (15). Briefly, frozen half brains were sonicated in urea lysis buffer, sonicated, centrifuged, reduced with DTT, and alkylated with iodoacetamide. 15 mg protein from each sample was digested with trypsin and purified over C18 columns for enrichment using phosphotyrosine pY-1000 motif antibodies #8803 (Cell Signaling, Danvers, Mass). Enriched peptides were purified over C18 STAGE tips (16) and subjected to LC-MS/MS analyses. Replicate injections of each sample were run non-sequentially. Phosphopeptides were eluted using 90-minute linear gradients of acetonitrile in 0.125% formic acid delivered at 280 nL/min. Tandem mass spectra were collected in a data-dependent manner using a Thermo Orbitrap Fusion™ Lumos™ Tribrid™ mass spectrometer, a top-twenty MS/MS method, a dynamic repeat count of one and a repeat duration of 30 sec. Real time recalibration of mass error was performed using lock mass (17) with a singly charged polysiloxane ion m/z = 371.101237.

MS/MS spectra were evaluated using Comet and the Core platform (17–19). Files were searched against the SwissProt *Mus musculus* FASTA database. A mass accuracy of +/-5 ppm was used for precursor ions and 0.02 daltons for product ions. At least one tryptic (K- or R-containing) terminus was required per peptide and up to four mis-cleavages allowed. Cysteine carboxamidomethylation was specified as a static modification. Methionine oxidation and phosphorylation on serine, threonine and/ or tyrosine residues were allowed as variable modifications. Reverse decoy databases were included for all searches to estimate false discovery rates (FDR) and filtered using Core’s linear discriminant module with a 1.0% FDR. Peptides were filtered for the presence of a tyrosine phosphorylated residue (strict motif) or serine/threonine phosphorylated residue within 2 amino acids of a tyrosine (lax motif). Quantitative results were generated using Skyline (20) to extract the integrated peak areas of the corresponding peptide assignments. Accuracy of quantitative data was ensured by manual review in Skyline and/or in the ion chromatogram files.

### Phosphatase/dephosphorylation assays

Recombinant PTPRD phosphatase protein (> 95% purity) was produced in *E Coli* from His-tagged constructs as described (21)(22). We synthesized human actinβ_1 KCDVDIRKDL[pY]ANTVLSGGTT, actinβ_2 IVRDIKEKLC[pY]VALDFEQEMA, actinβ_3 GDGVTHTVPI[pY]EGYALPHAIL, cofilin 1 GDVGQTVDDP[pY]ATFVKMLPDK and dock4 LGLDLVPRKE[pY]AMVDPEDISI phosphopeptides and compared these data to a positive control END[pY]INASL (Promega) phosphopeptide studied in the same experiments.

Orthophosphate release assays (Promega V2471) used Malachite green and molybdate with spectrophotometric detection of liberated free orthophosphate from test phosphopeptides compared to control/comparison phosphopeptides with assessments for the times indicated. Reactions were carried out in half-area 96-well plates with three wells dedicated for each time point. To each experimental well, we added a mixture of 18 μL of ultrapure water, 25 μL of running buffer (43.4 μM HEPES (pH 7.4), 2.2 mM dithiothreitol, 0.44% acetylated bovine serum albumin, 22.2 mM NaCl, 4.4 mM EDTA) and 1 μL of a 10mM DMSO solution of the desired peptide. 50 μL of molybdate dye mixture was added at t =0, followed by 5 μL of a 1:100 dilution of enzyme in dilution buffer (22.9 mM pH 7.4 HEPES, 1% acetylated bovine serum albumin, 4.6 mM dithiothreitol). In control experiments, we added enzyme that was heated to 100C for 20 min, cooled and added to reactions similarly. Other wells were initiated *via* the addition of 5 μL of the diluted enzyme mixture @ t=0 and terminated at the desired timepoints by addition of 50 μL of the dye solution. Wells were read @ 605 nm using a Spectromax spectrophotometer (Molecular Devices, San Jose CA).

### Data analyses

We used data from the phosphopeptides whose abundance was most different and statistically-significant (nominal) to identify genes presented tables in the body of the paper, for gene-based analyses of overlap with other datasets (using hypergeometric tests) and for tests of overrepresentation among Gene ontology terms. We used 1.5 or 2 fold cutoffs for genes in tables and p < 10^−8^ or < 10^−7^ for gene ontology terms following manual inspection of the datasets. Ambiguities in phosphopeptide assignment to genes are maintained in these tables.

Corresponding phosphopeptide-level datasets are included in the Supplement. Coexpression data come from Allen brain institute mouse cortex/hippocampus single cell RNA seq datasets.

## Results

### Phosphopeptides identified

We identified 1835 discrete phosphopeptides corresponding to 813 genes or groups of genes that were immunoprecipitated using phosphotyrosine antibodies and identified by these approaches.

### Phosphotyrosine phosphopeptides whose abundance increased with reduced PTPRD expression in the main dataset from young mice

We identified 101 phosphopeptides from 76 proteins (or groups of related proteins sharing the same sequence) that displayed > 1.5-fold increased abundance and nominal p < 0.05 in 2 month old PTPRD knockout *vs* wildtype mice from our main dataset (Table I; Supplement Table 1).

**Table I:**
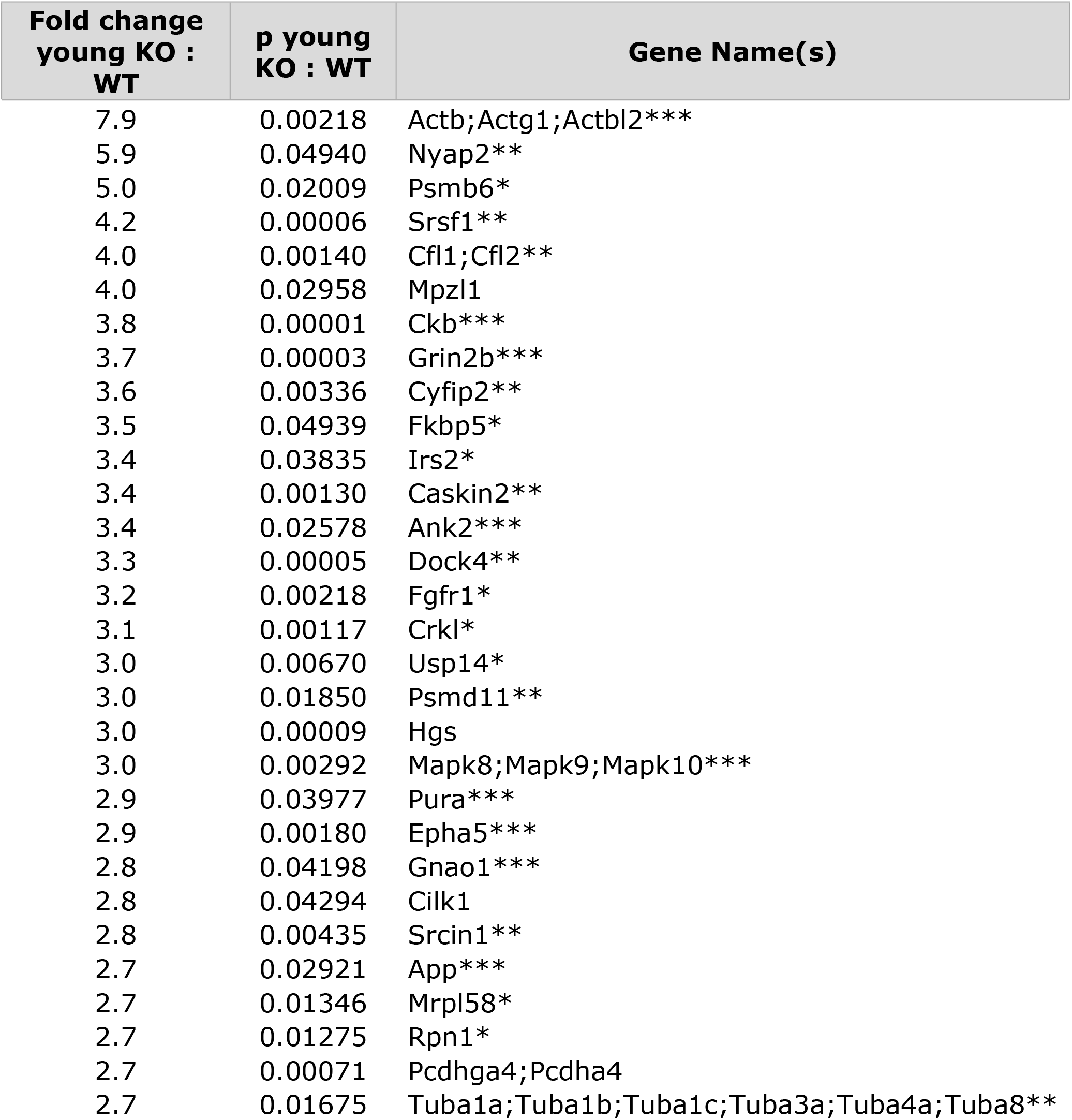
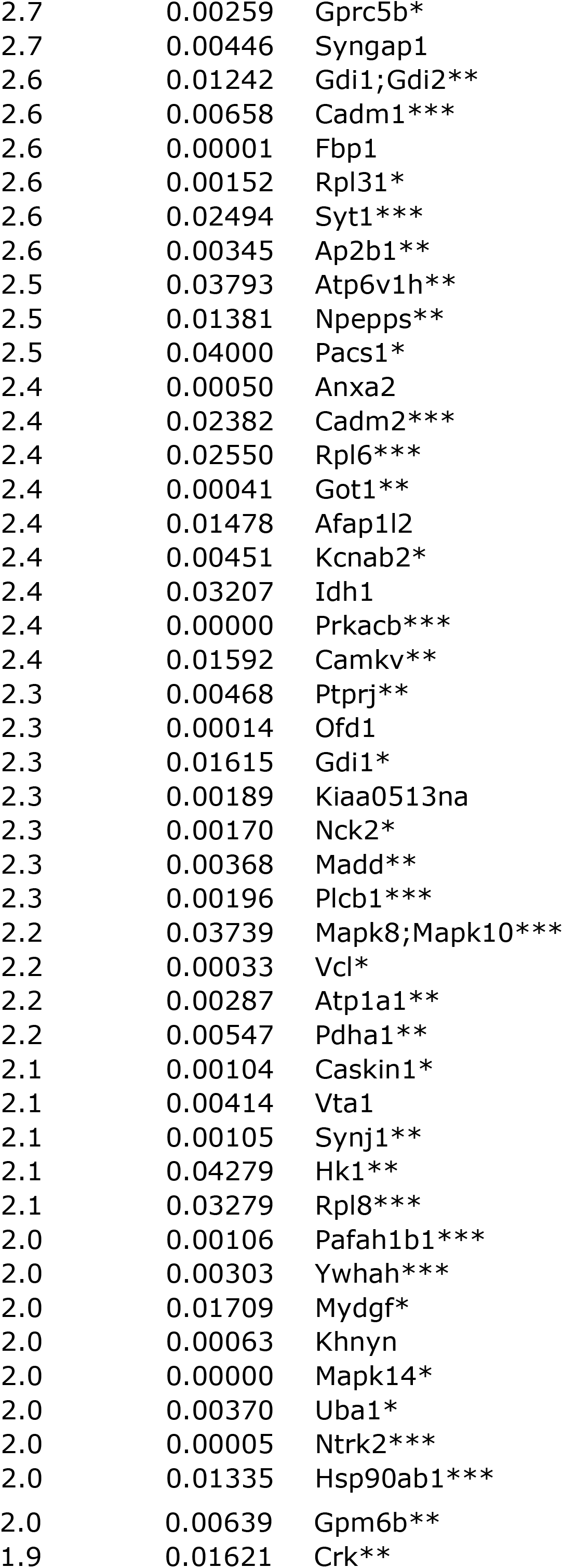
Genes encoding phosphopeptides with increased abundance in *2* mo old PTPRD knockout mice. Table 1: Genes containing phosphotyrosine phosphopeptides whose abundance is increased > 1.5 fold (fold increase column 1) with p < 0.05 (nominal p value column 2) in proteins extracted from brains of 2 month old PTPRD knockout mice compared to wildtype littermates. n = 4/genotype. Some phosphopeptides could come from several genes, as listed. Asterix number: relative extent of coexpression of the gene in the cell types that express PTPRD in mouse cerebral cortex + hippocampus RNAseq data (Allen brain institute). *** substantial coexpression; ** moderate coexpression; *coexpression in several cell types.

Gene ontology (GO) annotation of this list of genes documents overrepresentation of interesting GO terms with FDR-corrected p < 10^−4^, including: cell junction, synapse, cytoplasm, cell projection, postsynaptic density, asymmetric synapse, neuron projection, postsynaptic specialization, neuron to neuron synapse and glutamatergic synapse. (Supplement Table 2). Most of these genes are coexpressed with PTPRD in mouse cortical and hippocampal neuronal types defined by Allen Brain Institute single cell RNAseq datasets (Table 1). Only Caskin 2, Pcda4, Tub1a, Tub1c, Tub 3a, FBP1, Idh 1 and vta 1 fail to display substantial coexpression in at least one of the PTPRD-expressing neuronal types described in this data.

The group of coexpressed genes is thus likely to contain many PTPRD substrates, though some could also contain phosphotyrosines whose differential abundance could provide indirect adaptations to reduced PTPRD expression.

### Phosphotyrosine phosphopeptides whose abundance decreases with reduced PTPRD expression in the main dataset from young mice

We identified 101 phosphopeptides from 40 proteins (or groups of related proteins sharing the same sequence) that displayed > 1.5-fold decreased abundance and nominal p < 0.05 in 2 month old PTPRD knockout *vs* wildtype mice (Table II, Supplement Table 3).

**Table II:**
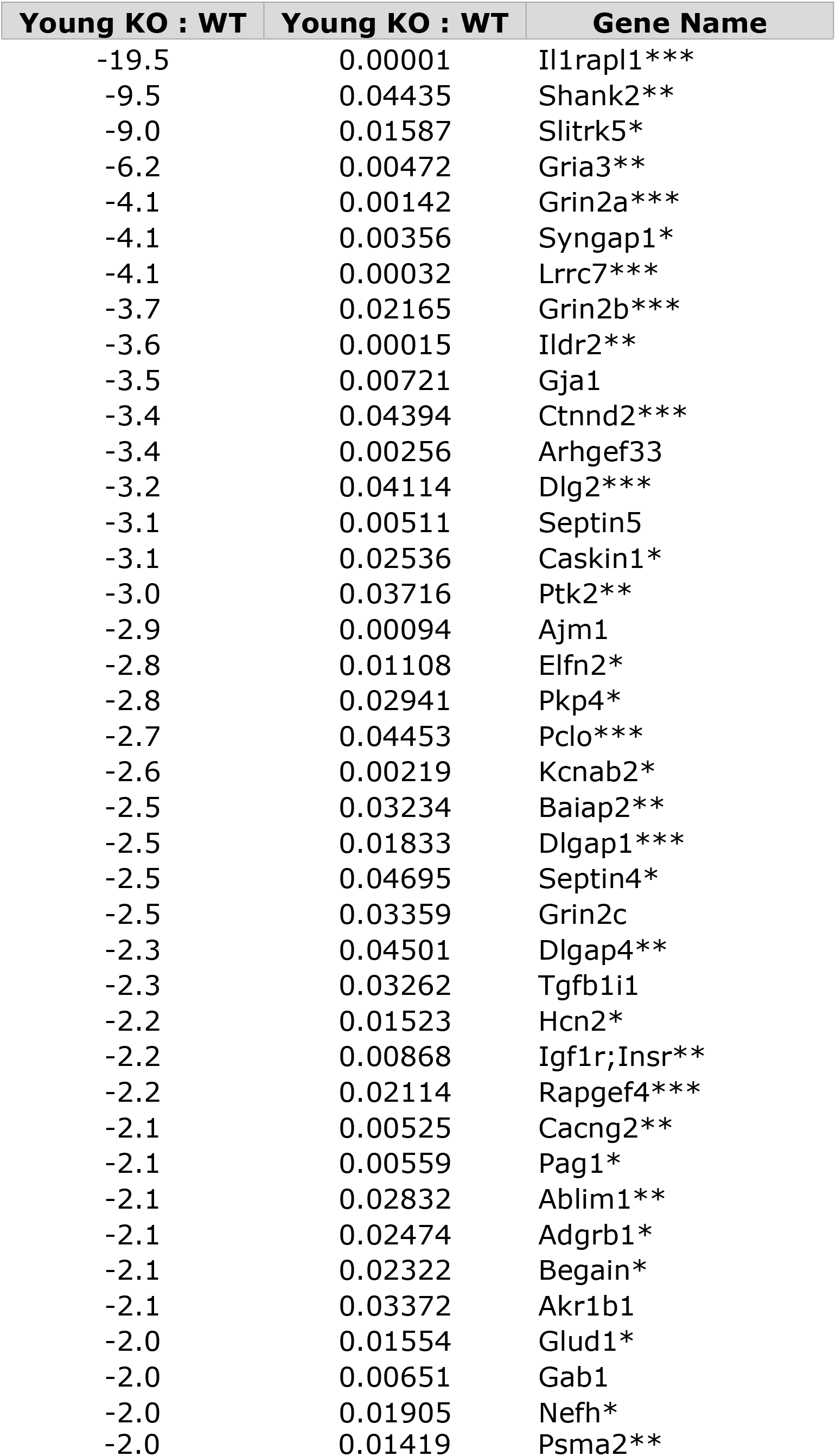
Genes encoding phosphopeptides with decreased abundance in 2 mo old PTPRD knockout mice. Table 2: Genes containing phosphotyrosine phosphopeptides whose abundance is decreased > 1.5 fold (fold decrease column 1) with p < 0.05 (nominal p value column 2) in proteins extracted from brains of 2 month old PTPRD knockout mice compared to wildtype littermates. n = 3 - 4/genotype. Some phosphopeptides could come from several genes, as listed. Asterix number: relative extent of coexpression of the gene in the cell types that express PTPRD in mouse cerebral cortex + hippocampus RNAseq data (Allen brain institute). *** substantial coexpression; ** moderate coexpression; *coexpression in several cell types

Gene ontology terms that are identified by this group of genes with FDR corrected p < 10^−4^ include: cell junction, postsynapse, postsynaptic specialization, synapse, postsynaptic density, asymmetric synapse, glutamatergic synapse, somatodendritic compartment, dendrite, neuron projection, ionotropic glutamate receptor complex, neurotransmitter receptor complex, plasma membrane region, ion channel complex and postsynaptic membrane (Supplementary Table 4).

These terms align with the idea that many of the adaptations to reduced PTPRD expression can be found in neuronal specializations that can be postsynaptic to often-presynaptically-expressed PTPRD (14). In particular, some of the most striking changes are found in candidate PTPRD ligands. The largest reduction in phosphorylation is in phosphopeptides in the PTPRD ligand Il1rapl1 (23). The third largest is in Slittrk5, another candidate PTPRD ligand (24). Each of these features is consistent with the idea that many of the reductions in protein tyrosine phosphorylation in PTPRD knockout mice could represent adaptations to PTPRD loss.

### Supporting studies of PTPRD phosphatase activities

We synthesized several actin as well as cofilin and dock4 phosphopeptides as candidate substrates for PTPRD’s phosphatase since these were: a) among those whose abundance increased most strikingly in two month old mice with reduced PTPRD expression; b) products of genes that are coexpressed abundantly with PTPRD in cerebral cortical neuronal cell types and c) products of genes that are good candidates for involvement in neuronal adaptations plausibly associated with PTPRD functions as a synaptic organizer (3, 25–28).

PTPRD phosphatase cleaved orthophosphate from each of these phosphopeptides and from our END(pY)INASL at rates much greater than those (essentially zero) for control experiments using boiled/inactive phosphatase. Rates for orthophosphate release ranged from almost twice those for the positive control END(pY)INASL for Dock4 and actinβ_3 phosphopeptides to about 20% of the positive control for the cofilin 1 phosphopeptide (Table 3). Each of these rates were significantly different from rates for experiments using boiled control phosphopeptide. These results support our phosphopeptide phosphoproteomic studies, since each of the tested candidate substrates is actively hydrolyzed by recombinant PTPRD phosphatase *in vitro.*

**Table III:** Relative rates of orthophosphate release by recombinant PTPRD phosphatase from phosphopeptides whose abundance increased in mice with reduced PTPRD expression. Table 3: Relative rates of orthophosphate release from synthetic phosphopeptides corresponding to candidate PTPRD substrates actin (three regions), cofilin and dock 4. Rates of orthophosphate release from phosphopeptides shown were compared to rates of release from positive control END(pY)INASL in the same experiment. Values are mean +/− SEM from three replicate experiments, each using triplicate assays and data from four time points.

|  | Ratio to pyEND |  |
| --- | --- | --- |
|  | Ratio | SEM |
| Actinβ_1 | 0.38 | 0.08 |
| Actinβ_2 | 0.71 | 0.07 |
| Actinβ_3 | 1.45 | 0.14 |
| Cofilin 1 | 0.17 | 0.02 |
| Dock4 | 1.88 | 0.14 |

### Phosphotyrosine phosphopeptides with smaller KO:WT differences in older vs younger mice

We identified 31 phosphopeptides from 24 genes that displayed > 1.5-fold smaller differences in abundance with p < 0.05 nominal significance in comparisons of old knockout to old wildtype *vs* young knockout to young wildtype mice (Table 4, Supplementary Table 5).

**Table IV:**
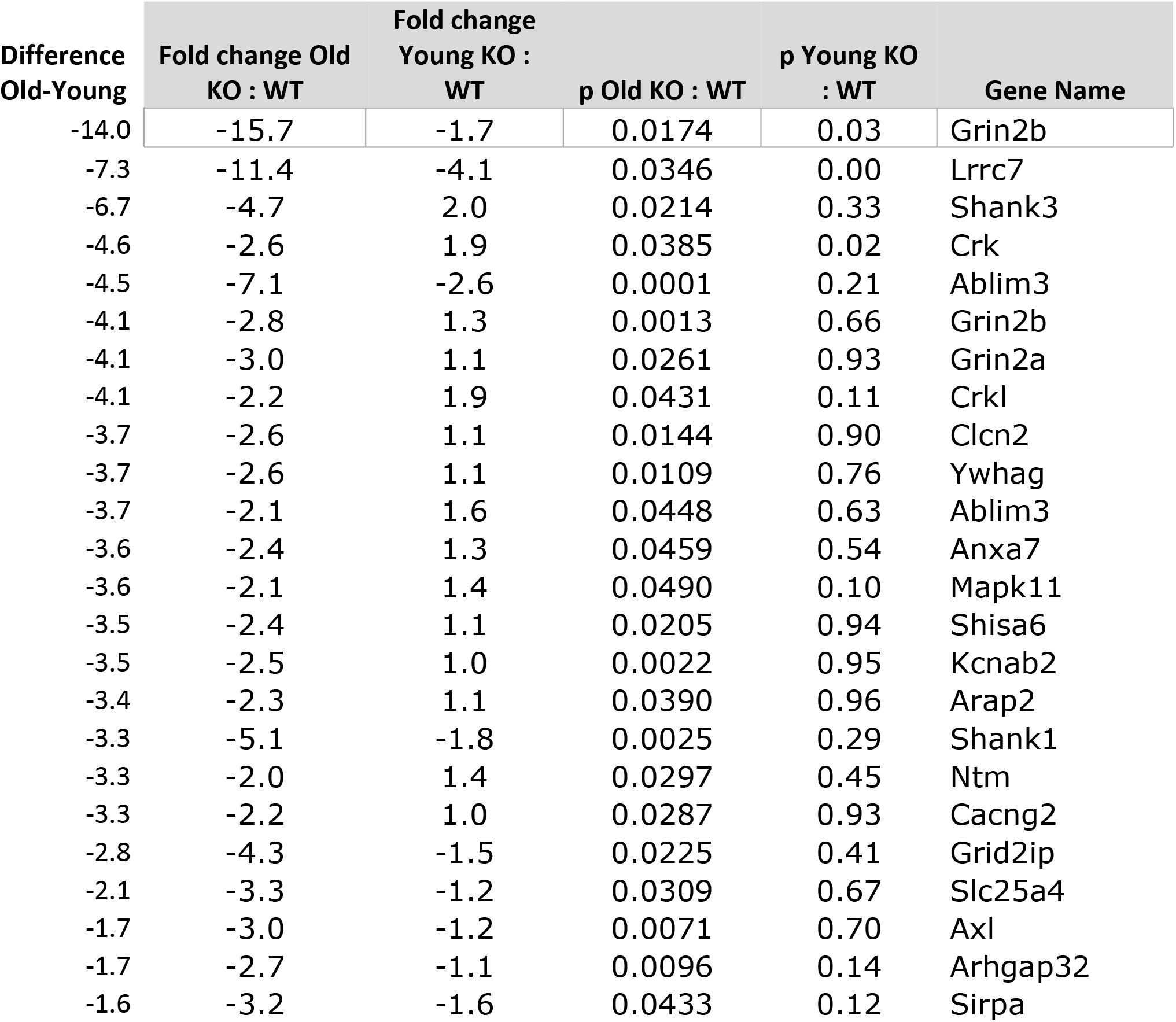
Genes encoding phosphopeptides with smaller KO:WT differences in 12 vs 2 mo old PTPRD knockout mice. Table 4: Genes containing phosphotyrosine phosphopeptides whose abundance change in 12 month old knockouts displays p < 0.05 and which display the greatest reductions in fold change compared to comparisons between 2 month old knockouts vs wildtype mice. n = 3 - 4/genotype. Some phosphopeptides could come from products of several genes, as listed. These genes represent candidates for interactions between effects of PTPRD expression and aging.

This group of genes provided p < 10^−7^ FDR-corrected significance for over-representation in GO categories: including neurotransmitter receptor complex, inotropic glutamate receptor complex, synapse, ion channel complex, transporter complex, postsynapse, postsynaptic density, asymmetric synapse, postsynaptic specialization, postsynaptic membrane and cell junction (Supplementary Table 6). These localizations fit a hypothesis that aging reduces the magnitudes of a number of the synaptic adaptations to loss of PTPRD.

### Phosphotyrosine phosphopeptides with larger KO:WT differences in older vs younger mice

We also identified 33 phosphopeptides from products of 24 genes whose abundance differences between knockout and wildtype mice were > 1.5-fold larger with p < 0.05 in comparisons of old knockout *vs* old wildtype mice (Table 5, Supplementary Table 7).

**Table 5:**
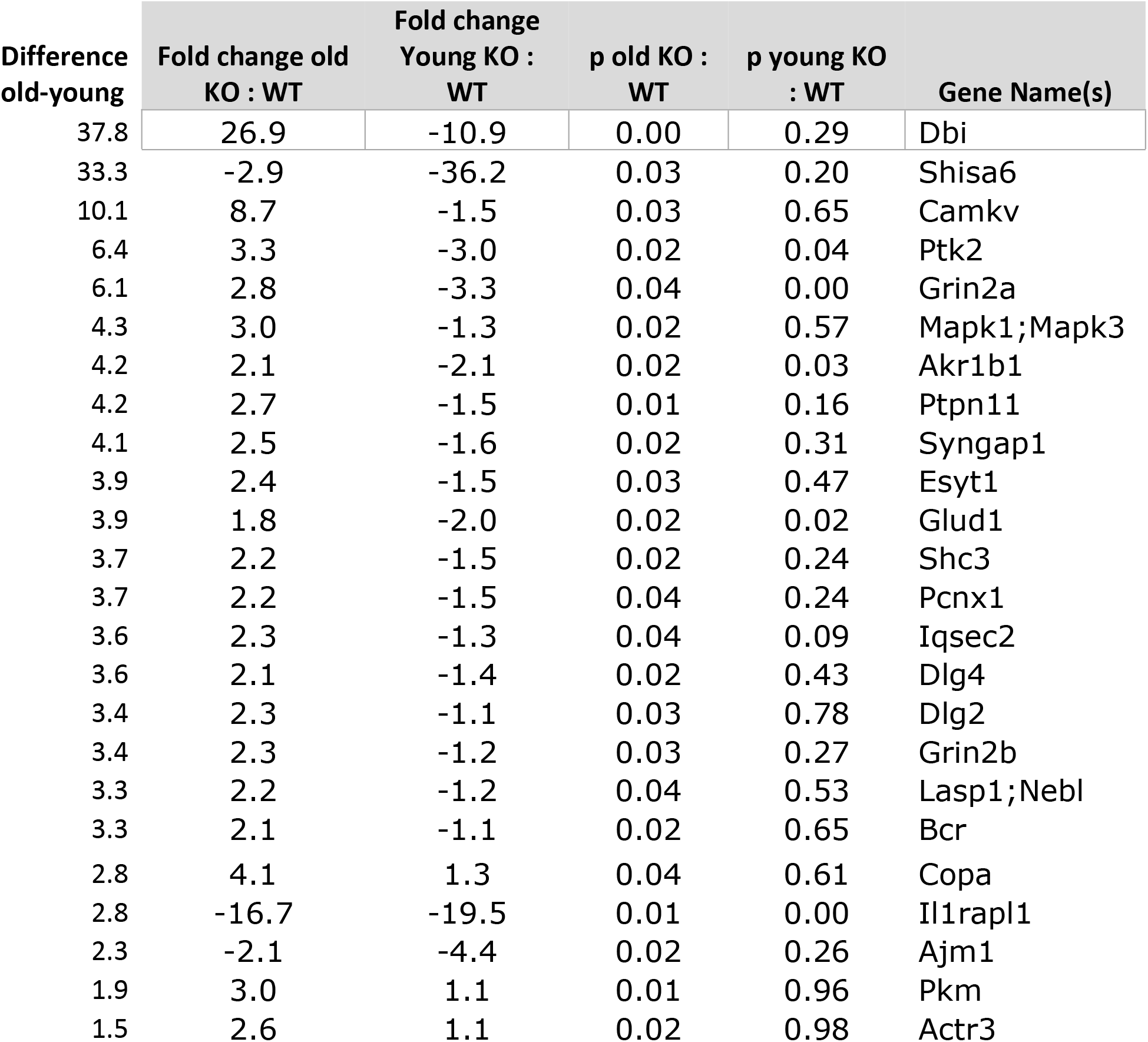
Genes containing phosphotyrosine phosphopeptides whose abundance change in 12 month old knockouts displays p < 0.05 and which display the greatest fold change compared to comparisons between 2 month old knockouts vs wildtype mice. n = 3 - 4/genotype. Some phosphopeptides could come from products of several genes, as listed. These genes represent candidates for interactions between effects of PTPRD expression and aging.

Overrepresented p < 10^−6^ GO terms supported aging influences on synaptic adaptations to loss of PTPRD including: glutamatergic synapse, cell junction, postsynaptic density membrane, synapse, postsynaptic specialization membrane, asymmetric synapse, postsynaptic density, neuron to neuron synapse, postsynaptic membrane, ionotropic glutamate receptor complex, intrinsic component of postsynaptic density membrane and neurotransmitter receptor complex (Supplementary Table 8). These localizations again fit with aging influences on the synaptic adaptations to loss of PTPRD.

### Replications in an independent dataset

We identified nominally-significant > 1.5 fold/ p < 0.05 changes in several of the top genes from our primary dataset in brains of independent groups of mice with a different brain freezing approach. Irs2, Kcnab2 and Pdha1 provided smaller magnitude, but p < 0.05 increased phosphopeptide abundance in this independent dataset. There was a trend (p = 0.08, hypergeometric test) toward significant overlap with the set of genes with more abundant phosphopeptide abundance in the data presented in Table I.

There was decreased expression (> 1.5-fold, p < 0.05) of Il1rapl1, Slitrk5 and Dlgap1in young homozygous knockouts in this dataset. The three overlapping genes whose phosphopeptide abundance decreased in young knockouts (Table 2) provided p = 10^−5^ (hypergeometric testing). Dlg2 and Grin2a were also decreased > 1.5 fold though with p > 0.05.

### Gene dose relationships in our independent dataset

Data comparing changes in homozygous *vs* heterozygous knockouts (*vs* wildtype mice) provide information about gene dose-response relationships for phosphopeptides from gene products that are most increased or decreased in abundance in knockouts. Phosphopeptides that displayed increased abundance in knockouts *vs* wildtype mice displayed typical gene dose-response patterns: heterozygotes displayed smaller differences from wildtype and less statistical significance than homozygotes for phosphopeptide products of each of the genes that displayed nominally-significant upregulation > 1.5-fold in our replication dataset.

Gene dose-response relationships for decreases were not as simple. Decreases in both IL1RAPL1 and SLITRK5 phosphopeptide abundance *vs* wildypype mice were smaller and did not reach nominal significance in heterozygotes, although the much larger declines found in homozygous knockouts reached high levels of significance, as noted above. By contrast, the declines in Dlg2, Dlgap1 and Grin2a phosphopeptide abundance were larger and reached higher levels of statistical significance for differences from wildtype in heterozygotes *vs* homozygotes.

## Discussion

The current results document differences in brain tyrosine phosphorylation that define groups of candidate PTPRD substrates, sets of candidate adaptations to loss of PTPRD activity and groups of potential aging effects on these candidate substrates and adaptations. We discuss the strengths and limitations of these results and the ways in which they point toward important roles for PTPRD activity. We note pathways whereby these changes could contribute to human and mouse model associations between PTPRD variation and addictions, RLS and Alzheimer’s disease pathophysiologies. We map the place that these brain phosphotyrosine phosphoproteomic results assume in the study of complex phosphorylation and dephosphorylation events in the brain.

The genes (or groups of genes) corresponding to phosphopeptides that display both > 1.5-fold increased abundance and nominal p < 0.05 in young PTPRD knockout *vs* wildtype mice form an interesting group. This group of genes contains many more whose products are expressed in presynaptic localizations (where much of PTPRD is expressed (14)) than expected by chance. Allen brain institute data supports colocalization of moderate to relatively high levels of expression of 45% of these genes in many of the neuronal subtypes that also express PTPRD (Table 1). Although it is possible that some of these increases in tyrosine phosphopeptide abundance come from adaptations to loss of PTPRD, this group of genes, overall, is likely to provide many candidate PTPRD substrates. There is significant, but imperfect overlap with other relevant datasets. There is significant overlap (11 genes, hypergeometric p = 10^−9^) between the gene products identified in our studies and the 59 genes/groups of genes whose tyrosine phosphorylation was reported increased by > 1.5 fold (p < 0.05) in studies of a different strain of PTPRD knockout mice that were reported while our work was in progress (14). There is also overlap with genes identified with increased tyrosine phosphorylation in *C. elegans* with deletion of the ptp-3 gene that corresponds to PTPRD, PTPRS and PTPRF (6 genes, hypergeometric p = 9 x 10^−5^) (13).

In other work we have identified 1.3-fold increases of abundance of a phosphotyrosine phosphopeptide with sequence shared between GSK3β and GSK3α in Western analyses of proteins from PTPRD knockout mice. This was one of the phosphopeptides increased in *C. elegans* ptp-3 knockouts (13). The more modest magnitude of our Western results could explain why it was not detected in our phosphoproteomic datasets with > 1.5-fold thresholds.

Actin, cofilin and dock 4 phosphopeptides identified as candidate substrates for PTPRD’s phosphatase in the phosphoproteomics datasets developed herein are each actual substrates for recombinant PTPRD phosphatase *in vitro.* Rates for orthophosphate release from these phosphopeptides range from brisk to very brisk, with release from the actinβ_3 peptide almost 50% greater and that from the dock4 phosphopeptide almost double the brisk rate of release from the active positive control END(pY)INASL. Each of the tested candidate substrates is thus actually hydrolyzed by recombinant PTPRD phosphatase *in vitro,* adding substantially to our overall confidence in the phosphoproteomic datasets.

There is also significant overall support for the set of phosphotyrosine phosphoproteins that display less abundance in 2 month old mice with reduced PTPRD expression. Allen brain institute data supports colocalization of moderate to relatively high levels of expression of 47% of these genes in many of the neuronal subtypes that also express PTPRD. There is significant overlap (8 genes, hypergeometric p < 10^−9^) between the gene products identified in our studies and the 43 genes/groups of genes whose tyrosine phosphorylation was reported decreased by > 1.5 fold (p < 0.05) in studies of a different strain of PTPRD knockout mice reported while our work was in progress (14). There is thus significant support for the validity of our set of downregulated genes as well as the GO links to postsynaptic locations that fit with adaptive roles for these reductions in phosphotyrosine phosphoprotein abundance. While several of these phosphopeptides have functional annotations in one of the best phosphopeptide annotation databases, future studies will be required to identify roles for most of these phosphotyrosines in regulating function. Roles in regulating the activities of products of the IL1RAPL1 and SLITRK 4 and 5 genes seem especially merited due to their roles as likely PTPRD ligands. In future work, we will assess possible contributions of changes in overall IL1RAPL1, SLITRK4 and SLITRK5 gene expression to these changes in abundance of the corresponding phosphotyrosine phosphopeptides.

There is evidence for both attenuation and increases in the changes in noted in young mice when 12 month-old knockout data is compared to data from wildtype littermate mice aged in parallel in the same facility. The genes whose products display less phosphotyrosine phosphoprotein abundance and those whose products display more phosphotyrosine phosphoprotein abundance in the older mice each provide candidates for interactions between age and level of PTPRD expression. From this group, we thus have candidates to participate in aging interactions with levels of PTPRD expression. Aging x level of expression changes are, in turn, candidates to contribute to the accumulation of phosphotau immunoreactivity that we have observed in aged mice with reduced PTPRD expression. This group also provides candidates to participate in the interactions between aging and altered PTPRD expression that are implied by the human associations between Alzheimer’s disease neurofibrillary pathology and intron 10 PTPRD genomic markers that are near those that we have associated with levels of PTPRD expression (4, 5).

There are limitations of these data. Similar sequences surrounding phosphotyrosines provide ambiguity concerning which exact actin, tubulin, MAP kinase, CAM kinase, disc-large or other gene products actually display increased abundance in mice with reduced PTPRD expression. Our results do not separate differences in levels of expression of genes whose phosphotyrosine phosphopeptide abundance that we monitor here from differences in the extent of tyrosine phosphorylation of these proteins. The wealth of information about the existence of tyrosine phosphorylation sites and their phosphorylation patterns has grown much faster than our understanding of the physiological or regulatory roles that many of these tyrosine phosphorylations provide. The apparent lower sensitivity of our datasets using a different freezing method, and thus the lower overlap of these independent results with our primary dataset, underscores the exquisite sensitivity of these phosphoproteomic analyses to the speed of tissue freezing.

Phosphotyrosine phosphoproteomic methods have been used to aid identification of tyrosine phosphorylated phosphoproteins whose abundance is changed in cells or tissues with changes in activity of other tyrosine phosphatases including products of the PTPRB (29), PTPRG (30), PTPN11 (31, 32) and PTP4A3 (33) genes. However, the relatively novelty of this area is highlighted in a recent review (34) that observes that only 3% of the phosphorylation sites annotated even in one of the most-updated databases (PhosphoSitePlus (35)) have a corresponding experimentally-validated human kinase. Our work thus adds to a modest but growing number of approaches to identification of protein tyrosine phosphatase substrates and adaptive changes to altered tyrosine phosphatase activities using this approach.

Human association datasets and mouse model results have motivated us to identify the first PTPRD phosphatase inhibitor lead compound, 7-BIA(10), and to find that quercetin and related flavanols provide the first PTPRD positive allosteric modulators/lead compounds (36). Transient pharmacological PTPRD phosphatase inhibition with 7-BIA leads to increases in brain pYGSK3β and pYGSK3α immunoreactivity that are about 2/3 the magnitude of changes induced by chronic genetic reductions in PTPRD expression in heterozygous knockout mice (37). The current phosphoproteomic datasets from knockout mice can thus provide a template for studies seeking effects of transient pharmacological modulation of brain PTPRD activities in ways that could improve understanding of PTPRD-associated pathophysiological disease processes in addiction, RLS and neurofibrillary pathologies, This type of work can aid more precise targeting of improved therapeutics to these pathophysiological mechanisms.

## Acknowledgements

PTPRD knockout mice were generously provided by N Uetani and Y Iwakura. A Nelson and K Abell performed phosphoproteomics sample preparation. We received valuable advice and assistance from W Wang, T Prisinzano, F I Carroll, A Sulima, K Rice, J Adair and O Kovbasnjuk. We are grateful to support from NIDA and NIA U01DA047713 and supplement, (J Acri and R Klein, program officers) and the Biomedical Research Institute of New Mexico GRU and IH are coinventors of a patent submitted by the VA Office of Technology Transfer “Compounds for treating tauopathies and Restless Leg Syndrome and methods of using and screening for the same” that covers work with PTPRD positive allosteric modulators. GRU serves without compensation on the Medical and Scientific Advisory board of the Restless Leg Syndrome Foundation. MPS is an employee of Cell Signaling Technology, Inc, Danvers, Massachusetts which offers commercial phosphoproteomics services.

## Supplementary Table Legends

Supplementary Table 1: Phosphotyrosine phosphopeptides and related genes whose abundance is increased > 1.5 fold with p < 0.05 in proteins extracted from brains of 2 month old PTPRD knockout mice compared to wildtype littermates. n = 4/genotype. Some phosphopeptides could come from several genes, as listed.

Supplementary Table 2: Gene ontology (cellular component) terms overrepresented (p < 10^−4^) in the list of genes in Table 1.

Supplementary Table 3: Phosphotyrosine phosphopeptides and related genes whose abundance is decreased > 1.5 fold with p < 0.05 in proteins extracted from brains of 2 month old PTPRD knockout mice compared to wildtype littermates. n = 4/genotype. Some phosphopeptides could come from several genes, as listed.

Supplementary Table 4: Gene ontology (cellular component) terms overrepresented (p < 10^−4^) in the list of genes in Table 2.

Supplementary Table 5: Phosphotyrosine phosphopeptides and related genes whose abundance is decreased > 1.5 fold more in old knockout *vs* wildtpe comparisons than in young knockout vs wildtype comparisons (and which display p < 0.05 in old knockout/wildtype comparisons. n = 3 - 4/ genotype/age group. Some phosphopeptides could come from several genes, as listed.

Supplementary Table 6: Gene ontology (cellular component) terms overrepresented (p < 10^−5^) in the list of genes in Table 4.

Supplementary Table 7: Phosphotyrosine phosphopeptides and related genes whose abundance is increased > 1.5 fold more in old knockout *vs* wildtpe comparisons than in young knockout vs wildtype comparisons (and which display p < 0.05 in old knockout/wildtype comparisons. n = 3 - 4/ genotype/age group. Some phosphopeptides could come from several genes, as listed.

Supplementary Table 6: Gene ontology (cellular component) terms overrepresented (p < 2 x 10^−6^) in the list of genes in Table 5.

